# Padé SSA: A frequency domain method for estimating the dynamics of stochastic reaction networks

**DOI:** 10.1101/2022.03.31.486511

**Authors:** Ankit Gupta, Mustafa Khammash

**Affiliations:** Department of Biosystems Science and Engineering, ETH-Zürich

**Keywords:** Biomolecular reaction networks, stochastic networks, stochastic simulation, Monte Carlo methods

## Abstract

Dynamic analysis and control of living cells relies on mathematical representations of cellular processes that are themselves modelled as biomolecular reaction networks. Stochastic models for biomolecular reaction networks are commonly employed for analysing intracellular networks having constituent species with low-copy numbers. In such models, the main object of interest is the probability distribution of the state vector of molecular counts which evolves according to a set of ordinary differential equations (ODEs) called the Chemical Master Equation (CME). Typically this set is very large or even infinite, making the CME practically unsolvable in most cases. Hence the outputs based on the CME solution, like the statistical moments of various state components, are generally estimated with Monte Carlo (MC) procedures by simulating the underlying Markov chain with Gillespie’s Stochastic Simulation Algorithm (SSA). However to obtain statistical reliability of the MC estimators, often a large number of simulated trajectories are required, which imposes an exorbitant computational burden. The aim of this paper is to present a frequency domain method for mitigating this burden by exploiting a small number of simulated trajectories to robustly estimate certain *intrinsic eigenvalues* of the stochastic dynamics. This method enables reliable estimation of time-varying outputs of interest from a small number of sampled trajectories and this estimation can be carried out for several initial states without requiring additional simulations. We demonstrate our method with a couple of examples.

## I. Introduction

The paradigm of reaction networks presents a rich modelling framework for processes in many biological disciplines, such as, systems biology, epidemiology, pharmacology and ecology. It is known that deterministic formulations of the dynamics, based on ODEs, become inaccurate when the network contains species present in low copy-numbers. In such cases the reactions become discrete-events, making the dynamics intrinsically noisy, that can significantly affect the macroscopic of the system [1]. To incorporate this noise, stochastic reaction network models become necessary and in these models the dynamics is commonly represented by a continuous-time Markov Chains (CTMC) [2]. In the emerging field of *Cybergenetics* [3], which aims to design and engineer intracellular control networks in living cells, the stochastic modelling paradigm is becoming an indispensable analytical tool, as these controller circuits are naturally required to work in low copy-number regimes so as to impose a small metabolic burden on the host cell [4].

Suppose the stochastic reaction dynamics is given by CTMC (*X*(*t*))_*t*≥0_ where *X*(*t*) is the vector of molecular counts of all the species at time *t*. Let *p*(*t*) be the probability distribution of the random state *X*(*t*) at time *t*, i.e.

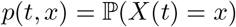

for each state *x* in the state-space ℰ of accessible states. The set of ODEs governing the dynamics of *p*(*t*) is called the CME (see (10)) and the number of ODEs in this set is the same as the number of states in ℰ which can be very large, and in most cases, infinite. Commonly, instead of the full CME solution *p*(*t*), one is interested in the expectation of some output function *f* : 𝒮 → ℝ under the probability distribution *p*(*t*), i.e.

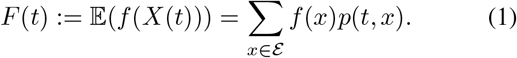

For example, if 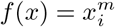, then the output of interest is the *m*-th moment of the molecular count of the *i*-th species at time *t*, i.e.

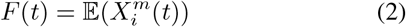

and when for some subset *A* ⊂ ℰ

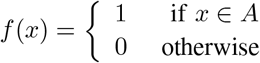

then the output is the probability of the state *X*(*t*) being in *A* at time *t*, i.e.

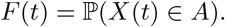

When the output is of the form (2), then in some cases its dynamics can be directly computed, without first solving the full CME. This is done by solving a system of ODEs for the moments [5]. However this ODE system may not be closed due to the *moment-closure* issue, and even though approximate techniques have been developed to circumvent this problem, the accuracy of these approaches cannot be ascertained in advance [6]. The standard approach for estimating the trajectory of *F* (*t*) is by simulating the CTMC via Gillespie’s Stochastic Simulation Algorithm (SSA) [7] and applying MC estimators to estimate the expectation at various time-points. Suppose we are interested in estimating the output dynamics in the time-period [0, *T*]. For this we simulate *N* CTMC trajectories (*X*^(*n*)^(*t*))_*t*≥0_ for *n* = 1, …, *N* and discretise the time-period as

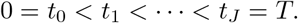

Then at each *t*_*j*_ the output *F* (*t*_*j*_) can be estimated by the standard MC estimator

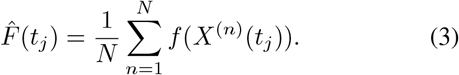

This estimator is unbiased and its statistical error, measured by its standard deviation, scales like 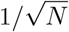. This scaling factor clearly shows how this estimator benefits from averaging w.r.t. population (i.e. trajectories) but this benefit accrues as 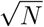 (population-size) even though the computational costs accrue like *N*. This mismatch highlights the difficulty in obtaining statistically reliable estimates, because simulating trajectories is computationally very expensive due to the time-step being inversely proportional to the fastest reaction’s timescale [7]. There are strategies to mitigate this problem and make simulations faster, e.g. employing *τ* -leaping methods [8] or exploiting timescale separation [9], but these strategies do not work in general and their accuracy cannot be easily checked.

In this paper, rather than finding efficient ways to generate stochastic trajectories, we adopt an alternate approach. We assume that we have a fixed computational budget to generate a small number of CTMC trajectories with SSA, and our aim is to exploit these trajectories in a better way than the standard approach in order to gain more information about the dynamics. In particular we work in the frequency domain and estimate the Laplace transform of the output trajectory, i.e.

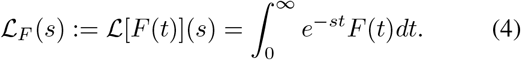

This problem is mathematically equivalent to the original problem of estimating *F*(*t*), and once we accurately estimate ℒ_*F*_(*s*) we can recover *F*(*t*) by applying the inverse Laplace transform. We shall develop our method under the assumption that the underlying CTMC is *exponentially ergodic*, i.e. starting from any initial state the solution of the CME relaxes to a unique stationary distribution, exponentially fast in time. This assumption is often satisfied by biological networks (see [10]) and it allows us to express ℒ_*F*_(*s*) as

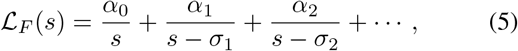

where *σ*_1_, *σ*_2_, … are *intrinsic eigenvalues* of the stochastic dynamics that do not depend on the output function *f* or the initial state *X*(0) = *x*_0_ of the CTMC trajectory (*X*(*t*))_*t*≥0_. The dependence on *x*_0_ and *f* is fully absorbed by the coefficients *α*_0_, *α*_1_, …. Expression (5) is equivalent to the following series expansion for *F*(*t*)

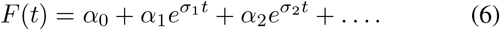

If we only keep the first (*q* + 1) terms in (5) then ℒ_*F*_(*s*) can be approximated as

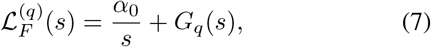

where *G*_*q*_(*s*) is a rational function whose denominator is a monic polynomial of degree *q* and the numerator is a polynomial of degree (*q* − 1).

Note that *α*_0_ is the steady-state output value, which due to ergodicity, coincides with the long-term time-average of the output for any CTMC trajectory (*X*(*t*))_*t*≥0_

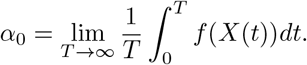

Hence if we have *N* such trajectories then *α*_0_ can be estimated as

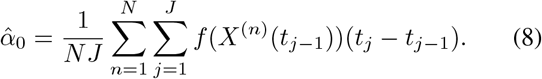

Notice that this estimator benefits from two layers of averaging - one is w.r.t. population (i.e. trajectories) as in the standard MC estimator (3), and the other is w.r.t. time *t*. This additional time-averaging improves the estimate *vis- à-vis* the standard MC estimator. Extending this idea further we develop such two-layered estimators for inferring the rational function *G*_*q*_(*s*) by computing the multipoint Padé approximant [11] of ℒ_*F*_(*s*). Henceforth we shall refer to our method as *Padé SSA* as it combines exact trajectory simulations obtained by SSA with Padé approximations. Such approximations have a rich history in control theory and applied mathematics and many works have shown their remarkable accuracy and convergence properties [12].

Observe that the denominator polynomial of *G*_*q*_(*s*) would have the intrinsic eigenvalues as roots and hence it would not depend on the initial state *X*(0) = *x*_0_ of the CTMC. Exploiting this fact we shall show how once this denominator polynomial has been estimated for some initial state *x*_0_, we can easily recompute the numerator polynomial of *G*_*q*_(*s*) for any initial state *x*_0_ *without needing to generate any additional CTMC trajectories*. This is a critical bottleneck of the standard approach because the CTMC trajectories generated with one initial state cannot be used to estimate the dynamics with a different initial state, essentially forcing the users to generate fresh CTMC trajectories for each initial state of interest.

Finally we would like to point out that our *Padé SSA* method is inspired by a similar method called *Padé PSD* that we have recently developed for the estimation of power spectral density (PSD) of stochastic single-cell trajectories from reaction network models (see [13]).

## II. Preliminaries

### A. Stochastic Reaction networks

We work with the standard CTMC model of a reaction network [2]. Suppose the network consists of *d* biomolecular species **X**_**1**_, …, **X**_**d**_, and *K* reactions with propensity functions *λ*_1_, …, *λ*_*K*_ and stoichiometric vectors *ζ*_1_, …, *ζ*_*K*_. The state of the system is the vector of copy-numbers of all the species *x* = (*x*_1_, …, *x*_*d*_). When the state is *x*, reaction *k* fires with rate *λ*_*k*_(*x*) and it displaces the state to *ζ*_*k*_. Formally, the CTMC (*X*(*t*))_*t*≥0_ representing a reaction dynamics can be specified by its generator (see Chapter 4 in [14]) which is the operator defined by

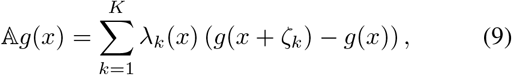

for any real-valued function *g* over the state-space ℰ. For any state *x* ∈ ℰ, the probability *p*(*t, x*) = ℙ (*X*(*t*) = *x*) evolves in time according to the Chemical Master Equation (CME)

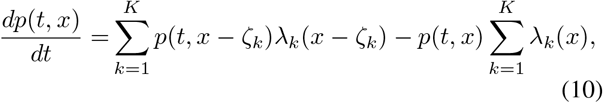

for each *x* ∈ ℰ. This system has as many ODEs as the number of states in ℰ which is typically very large or infinite. Hence the CME can rarely be directly solved and output quantities like (1) are estimated by simulating the CTMC with Gillespie’s SSA [7] or similar Monte Carlo procedures.

The CTMC is said to be ergodic if *p*(*t*) converges to a unique stationary distribution *π* as *t* → ∞, regardless of the initial state of the system. If this convergence is exponentially fast then the CTMC is called *exponentially ergodic* and this is what we assume in this paper. We have developed extensive computational procedures for checking exponential ergodicity for stochastic reaction network models (see [10] and [15] for more details).

### B. Frequency domain analysis

Corresponding to the CTMC (*X*(*t*))_*t*≥0_ with generator 𝔸 we define the resolvent operator as

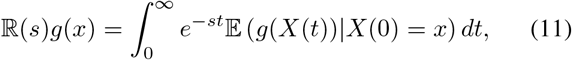

where *s* is the complex-valued frequency variable. In other words ℝ(*s*)*g*(*x*) is the Laplace transform of the map from time *t* to the conditional expectation 𝔼(*g*(*X*(*t*)) *X*(0) = *x*). Of course, for a given initial state *x*_0_ the Laplace transform ℒ_*F*_(*s*) is essentially the resolvent operator applied to the function *f* and evaluated at *x* = *x*_0_

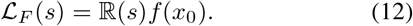

Note that *σ*_0_ = 0 is an eigenvalue of the generator 𝔸 and suppose *σ*_1_, *σ*_2_, … are the non-zero eigenvalues of 𝔸. These eigenvalues do not depend on *x*_0_ and *f* and we assume that all eigenvalues are distinct and arranged in descending order of their real parts (which are negative due to ergodicity). The eigen-decomposition of the resolvent operator (see the supplement of [13]) provides the expression (5), where each *α*_*j*_ captures the contribution in the dynamics of the eigenmode corresponding to *σ*_*j*_. Under the assumption of exponential ergodicity, the resolvent operator is compact and this ensures that it can be accurately approximated by a finite-rank operator [16] and hence we would have *α*_*j*_ → 0 as *j* → ∞. This suggests that (7) can be an accurate approximation of (5). In the case that eigenvalues are not distinct, the expression (5) would change but the argument for the accuracy of (7) remains valid.

## III. PadÉ SSA

We now describe our method Padé SSA for estimating ℒ_*F*_(*s*) from a handful of CTMC trajectories simulated with SSA over time-interval [0, *T*]. Specifically we shall estimate 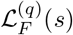 defined by (7). Note that the steady-state value *α*_0_ can be estimated as (8) and the main challenge is to identify the rational function

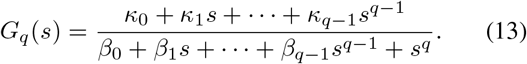

If we define functions 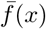 and 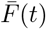 as

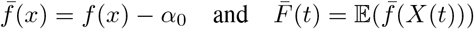

respectively, then the Laplace transforms ℒ_*F*_(*s*) and 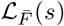 are related as

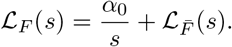

This shows that *G*_*q*_(*s*) must serve as an approximant to the Laplace transform 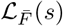. As these functions are complex-analytic it suffices to construct this approximation over the extended positive real-time (0, ∞]. We shall employ the method of multipoint Padé approximation [11] to identify the 2*q* coefficients (viz. *κ*_0_, …, *κ*_*q*−1_, *β*_0_, …, *β*_*q*−1_) by matching a certain number of derivatives *ρ*_1_, …, *ρ*_*L*_, at several arbitrarily chosen points *s*_1_, …, *s*_*L*_ in (0, ∞]. For this we need to find a way to estimate the higher-order derivatives of 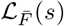 at various values of *s*.

Let us define the *m*-th Padé derivative of function 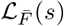 at *s* = *s*_*ℓ*_ *<* ∞ as

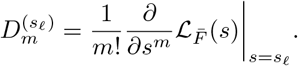

From (11) and (12) it can be seen that

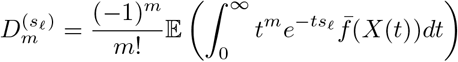

where (*X*(*t*))_*t*≥0_ is the CTMC with initial state *X*(0) = *x*_0_. By simulating *N* such CTMC trajectories we can estimate this Padé derivative as

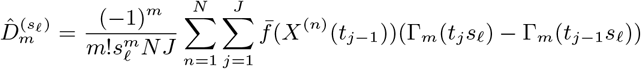

where Γ_*m*_(*z*) is the incomplete Gamma function given by

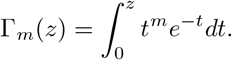

For the case *s*_*ℓ*_ = ∞ we shall define the *m*-th Padé derivative of function 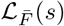 at *s* = ∞ as

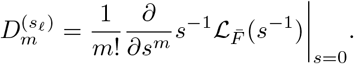

Note that

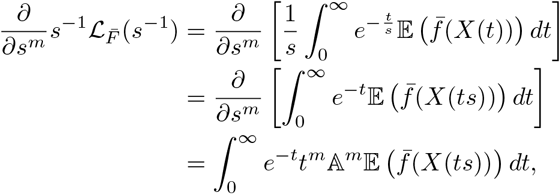

where the last relation follows from Dynkin’s formula [14]. Here 𝔸^*m*^ denotes the *m*-th iterate of the generator 𝔸 with 𝔸^0^ = **I** (the identity operator). Since *X*(0) = *x*_0_ and

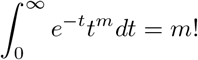

setting *s* = 0 in this expression shows that the *m*-th Padé derivative of function 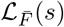 at *s* = ∞ is given by

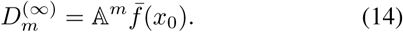

Observe that 𝔸^*m*^*f* (*x*_0_) can be easily computed recursively (see [13]) and so we do not need to estimate it with simulations, unlike the Padé derivatives for finite values of *s*.

Suppose for now that all the required Padé derivatives have been estimated or computed. Then by matching these derivatives with the rational function *G*_*q*_(*s*) given by (13) we can construct a linear system of the form

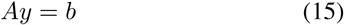

to solve for the vector of unknown coefficients *y* = (*κ*_0_, …, *κ*_*q*−1_, *β*_0_, …, *β*_*q*−1_). Here *A* is a matrix with dimensions *ρ*_sum_×2*q* and *b* is a vector with dimensions *ρ*_sum_×1, where

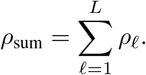

This system can be both underdetermined or overdetermined and so we solve it in least-squares sense. We add positivity constraints on the denominator coefficients *β*_0_, …, *β*_*q*−1_, which is necessary to ensure the stability of the dynamics. The exact expressions for *A* and *b* were derived in [13] and they can be found in the Appendix of this paper.

Now let us consider the scenario that the coefficient vector *y* = (*κ*_0_, …, *κ*_*q*−1_, *β*_0_, …, *β*_*q*−1_) has been estimated with some initial state *x*_0_ of the CTMC and we now want to estimate the output trajectories with another initial 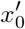. This can be done by matching the Padé derivatives at *s* = ∞, without the need for generating fresh CTMC trajectories with the new initial state. To see this, recall that the denominator coefficients *β*_0_, …, *β*_*q*−1_ will not depend on the initial state and the same holds for the stead-steady value *α*_0_ due to ergodicity. Moreover we can easily compute the Padé derivatives 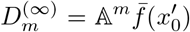 at the new initial state. Then the expressions for *A* and *b* in the linear system imply that the new numerator coefficients *κ*_0_, …, *κ*_*q*−1_ can be computed as

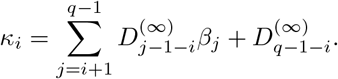

## IV. Examples

It is well-known that there are two fundamental topologies that exhibit robust perfect adaptation to input perturbation [17]. These topologies are known as *Incoherent Feedforward Loop (IFFL)* and *Negative Feedback Network (NFB)*. We demonstrate our Padé SSA method using simple three-node examples of these topologies. In both the examples when we apply Padé SSA we set *L* = 4, *s*_1_ = 0.25, *s*_2_ = 0.5, *s*_3_ = 1 and *s*_4_ = ∞ and *ρ*_1_ = *ρ*_2_ = *ρ*_3_ = *ρ*_4_ = 1.

### A. Incoherent Feedforward Loop (IFFL)

The IFFL network (see Figure 1(A)) consists of the following reactions

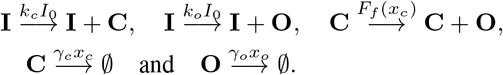

**Fig. 1.**
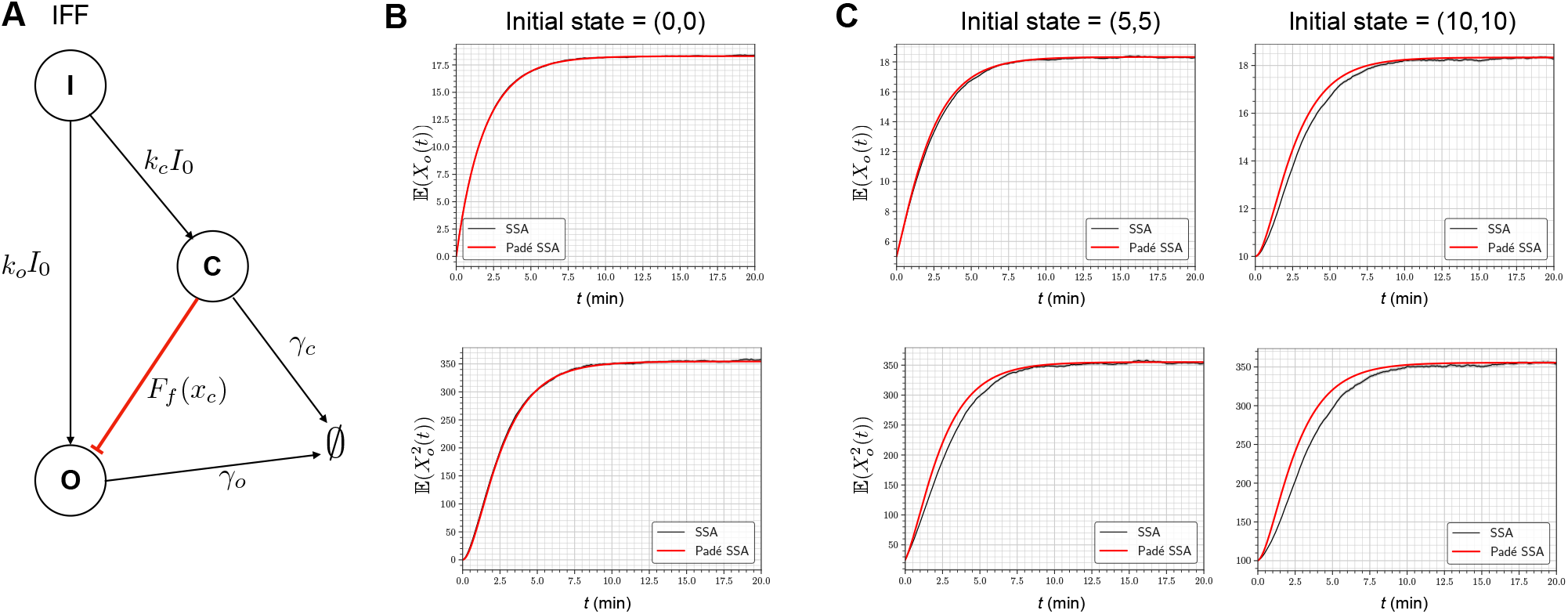
(A) Depicts the IFFL network where the input **I** produces the output **O** and the controller **C**. The controller then represses the production of the output. (B) Comparison of the dynamics of the first two moments estimated by the standard SSA approach and our Padé SSA method. (C) Moment dynamics estimated by our Padé SSA method for a couple of initial states. For Padé SSA the same set of trajectories was used for estimation for all the initial states. On our computer, generating *N* = 10^4^ trajectories with SSA took 39 minutes on average. Given the trajectories, the computational time for Padé SSA is less than 0.1 seconds.

The reaction propensities are stated above the reaction arrows, and *x*_*c*_ and *x*_*o*_ denote the copy-numbers of the controller species **C** and the output species **O**. The input abundance level *I*_0_ is constant and the repression of the production of **O** by **C** is given by a monotonically decreasing function *F*_*f*_ (*x*_*c*_). For our analysis we choose *F*_*f*_ (*x*_*c*_) as the nonlinear Hill function

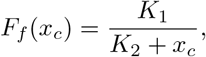

where *K*_1_, *K*_2_ are positive constants.

We shall estimate the dynamics of the first two moments 𝔼(*X*_*o*_(*t*)) and 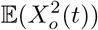 of the output species copy-numbers with the following parameter values

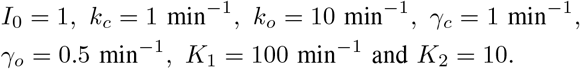

We simulate *N* = 10^4^ SSA trajectories in the interval [0, *T*] with *T* = 100 and in Figure 1(B) we compare the estimates we obtain with our Padé SSA method to the standard SSA approach. Observe that the standard SSA estimator is quite noisy while Padé SSA provides a smooth estimate of the dynamics which agrees well with it. In Figure 1(C) we re-estimate the dynamics with Padé SSA for a couple of initial states and compare the results with the SSA estimator. Note that no new SSA trajectories were generated in this analysis unlike the standard SSA approach where fresh trajectories need to be generated for each new initial state.

### B. Negative Feedback Network (NFB)

The NFB network (see Figure 2(A)) consists of the following reactions

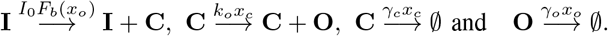

**Fig. 2.**
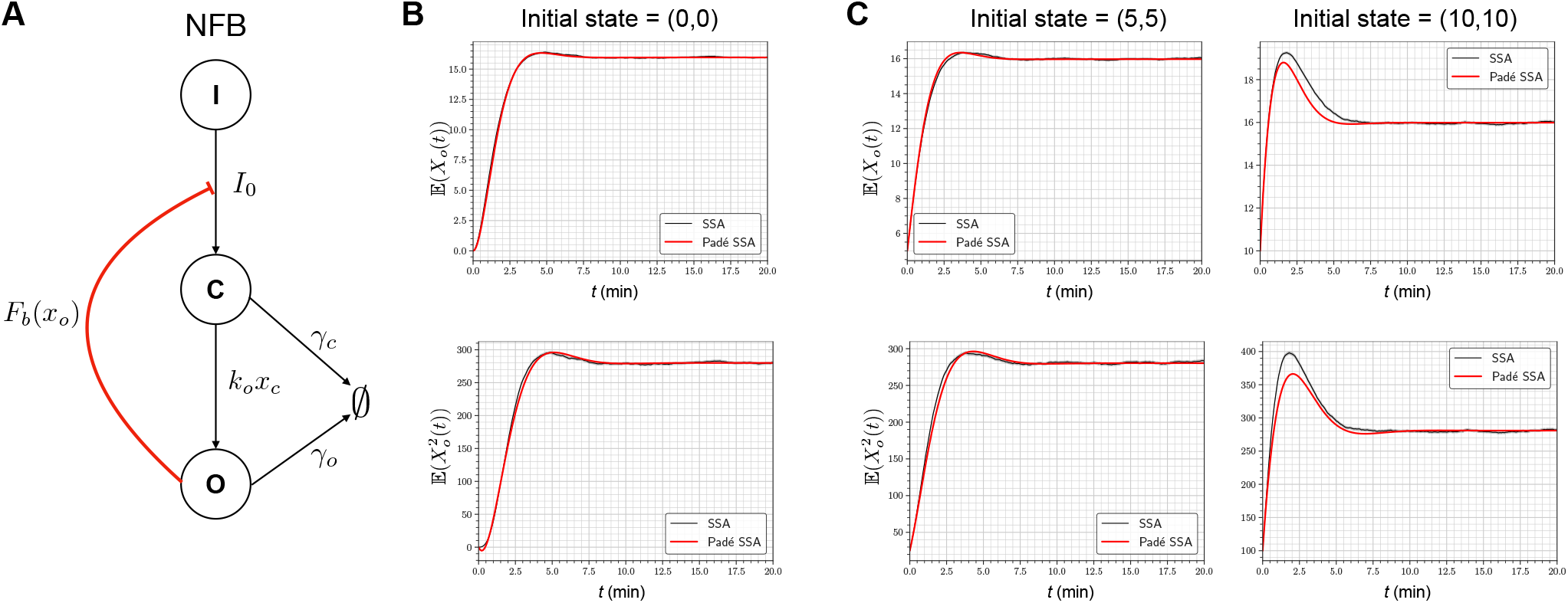
(A) Depicts NFB network where the input **I** produces the controller **C** which in turn produces the output **O** that then inhibits the production of **C** from **I**. (B) Comparison of the dynamics of the first two moments estimated by the standard SSA approach and our Padé SSA method. (C) Moment dynamics estimated by our Padé SSA method for a couple of initial states. For Padé SSA the same set of trajectories was used for estimation for all initial states. On our computer, generating *N* = 10^4^ trajectories with SSA took 43 minutes on average. Given the trajectories, the computational time for Padé SSA is less than 0.1 seconds.

In this example the monotonically decreasing function *F*_*b*_(*x*_*o*_) denotes the repression of the production of the controller species **C** by the output species **O**. We shall choose this feedback function as the nonlinear Hill function

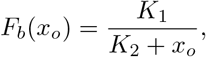

where *K*_1_, *K*_2_ are positive constants. As before we shall estimate the dynamics of the first two moments 𝔼(*X*_*o*_(*t*)) and 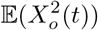 with the following parameter values

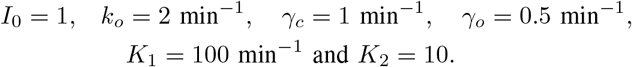

We simulate *N* = 10^4^ SSA trajectories in the interval [0, *T*] with *T* = 100 and Figure 2(B) provides a comparison of the estimated dynamics between Padé SSA and the standard SSA approach. Figure 2(C) we re-estimate the dynamics with Padé SSA for a couple of initial states and compare the results with the SSA estimator.

## V. Conclusion

The Chemical Master Equation (CME) for stochastic reaction networks is notoriously difficult to solve due to an inherent curse-of-dimensionality. Commonly, dynamic outputs based on the CME solution, like means and variances, are estimated by simulating the underlying Markov chain with SSA and then applying standard Monte Carlo estimators. The main problem with this approach is that often a large sample-size (i.e. number of trajectories) is required to attain statistical reliability, which can be computationally difficult to realise in practice. In this paper our goal is to mitigate this issue by estimating the output dynamics in the frequency domain through the use of Laplace transforms. We argue that the Laplace transform of the desired output trajectory can be accurately described as a certain type of rational function of the frequency variable, and we demonstrate how this rational function can be identified using the theory of Padé approximations. This method, which we call *Padé SSA*, not only produces a statistically accurate estimate of the dynamics with a small number of trajectories, it also allows one to change the initial state and re-estimate the dynamics without the need to generate new stochastic trajectories. Hence the same set of trajectories provide estimates of the output trajectories for any initial state, unlike the standard SSA-based approach. We illustrate Padé SSA with a couple of examples.

## Appendix

The matrix *A* and vector *b* in linear system (15) are constructed by vertically stacking *A*^(*ℓ*)^ and *b*^(*ℓ*)^ for each *ℓ* = 1, …, *L*. Here *A*^(*ℓ*)^ is a *ρ*_*ℓ*_ × 2*q* matrix and *b*^(*ℓ*)^ is a *ρ*_*ℓ*_ × 1 vector. Their components in the case *s*_*ℓ*_ *<* ∞ are given by

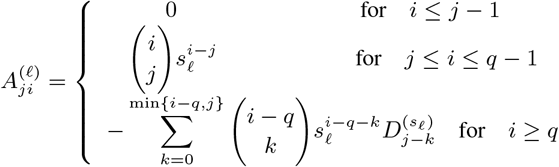

and 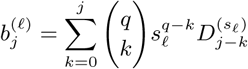.

In the case *s*_*ℓ*_ = ∞ these components are given by

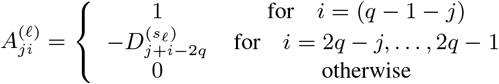

and 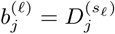.

